# Somatic Genetic Aberrations in Benign Breast Disease and the Risk of Subsequent Breast Cancer

**DOI:** 10.1101/613505

**Authors:** Zexian Zeng, Andy Vo, Xiaoyu Li, Ali Shidfar, Paulette Saldana, Xiaoling Xuei, Yuan Luo, Seema A. Khan, Susan E. Clare

## Abstract

It is largely unknown how the risk of development of breast cancer is transduced by somatic genetic alterations. To address this lacuna of knowledge and acknowledging that benign breast disease (BBD) is an established risk factor for breast cancer, we established a case-control study: The Benign Breast & Cancer Risk (BBCAR) Study. Cases are women with BBD who developed subsequent invasive breast cancer (IBC) at least 3 years after the biopsy and controls are women with BBD who did not develop IBC (median follow-up 16.6 years). We selected 135 cases and individually matched controls (1:2) to cases based on age and type of benign disease: non-proliferative or proliferation without atypia. Whole exome sequencing was performed on DNA from the benign lesions and from subsets with available germline DNA or tumor DNA. Although the number of cases and controls with copy number variation data is limited, several amplifications and deletions are exclusive to the cases. In addition to two known mutational signatures, a novel signature was identified that is significantly (p=0.007) associated with triple negative breast cancer. The somatic mutation rate in benign lesions is similar to that of invasive breast cancer and does not differ between cases and controls. Two mutated genes are significantly associated with time to the diagnosis of breast cancer, and mutations shared between the benign biopsy tissue and the breast malignancy for the ten cases for which we had matched pairs were identified. BBD tissue is a rich source of clues to breast oncogenesis.

**One Sentence Summary:** Genetic aberrations in benign breast lesions distinguish breast cancer cases from controls and predict cancer risk.

## Introduction

From 1989 to 2016 the mortality rate for breast cancer in the United States decreased by 40%^1^, a testament to the efficacy of targeted therapies, as well as to combinations and schedules of chemotherapeutics. During this same period breast cancer incidence rates remained static^1^; evidence of both our inability to implement existing prevention strategies, and to the paucity of novel, effective prevention strategies that target specific molecular risk pathways. Opportunities for prevention fall into three broad categories: surgical risk reduction, which is reserved for the small minority of women with high penetrance germline mutations^2^; lifestyle modification^3^, which is applicable to all, but difficult to implement; and medical prevention, which is of proven benefit to a broad swathe of intermediate and high risk women. Medical intervention appeared to have gained significant momentum with the publication of the P1 trial in 1998, which demonstrated a halving of breast cancer incidence with use of the prototypic prevention drug, tamoxifen^4^. However, despite level I evidence of efficacy of several selective estrogen receptor modulators (SERMs)^4–9^, and two aromatase inhibitors (AIs)^10,11^; the uptake of pharmacological prevention has languished, with a recent meta-analysis documenting acceptance of proven therapies by only 16% of eligible women^12^, a problem further compounded by poor adherence by those who initiate treatment^13^. This dismal uptake rate means that chemoprevention as practiced today will prevent fewer than 1% of breast cancers^14^. Major barriers are two-fold: the reluctance of healthy women to accept drugs for a disease that they may or may not experience in the future; and unwillingness of the same healthy women to experience side effects that impair quality of life and may compromise health^15^.

Almost 30 years after the first publication of the Gail model^16^ breast cancer risk stratification remains imprecise and insensitive to breast cancer subtype. In an analysis of data from the Womens’ Health Initiative, the Gail Model displayed modest ability to predict the risk of breast cancer (AUC = 0.58, 95% CI = 0.56 to 0.60)^17^. Among women at high risk of breast cancer, for example those diagnosed with atypical hyperplasia, neither the Gail model/Breast Cancer Risk Assessment Tool^18^ nor the Tyrer-Cuzick model^19^ performed well. This is a significant barrier to implementation of established medical interventions for disease prevention, and to the development of new, targeted intervention strategies for women at risk. If prevention strategies are to have an impact of reducing breast cancer incidence, biomarkers of substantially elevated breast cancer risk must be identified and validated; and it must be determined how breast cancer risk is transduced at the molecular level. Only then will it be possible to develop targeted interventions that eliminate or mitigate breast cancer risk.

Benign breast disease is an established risk factor for breast cancer^20,21^ and about 30% of breast cancer cases have a history of benign breast disease^22^. Of the 1.7 million breast biopsies each year in the U.S.^23^, about 75% of these return a diagnosis of benign breast disease including atypical hyperplasia^22^. Given the disappointing results of risk models based on demographic and clinical data, we sought to evaluate the genetic landscape benign breast biopsies, and identify patterns which may presage the onset of malignancy. Starting in the embryo^24^, tissues accumulate DNA mutations over time^25^. Most of the mutations are repaired, many are inconsequential, but a few may lead to or cause cancer^26,27^. Before there is any histologic evidence of invasive cancer, histologically normal tissue and premalignant lesions contain molecular aberrations that are associated with malignancy^28,29^. For example, sun-exposed, physiologically normal eyelid skin has been shown to have a mutation burden of 2-6 mutations/megabase/cell, a rate similar to that observed in many cancers^30^. The processes that causes these mutations leaves an imprint on the genome^31^. In the sun-expose eyelid epidermis mutations occur within a pattern that mimics the Welcome Trust Sanger Institute (WTSI) Mutation Signature 7, which is associated with ultraviolet exposure and its consequent CC>TT dinucleotide mutations at dipyrimidines^32^. Exogenous or endogenous mutational process, such as that that produced WTSI signature 7, are chemical reactions with DNA. Where they occur within a sequence is determined by various physicochemical constraints. While mutational processes are responsible for the creation of mutations, these mutations are selected for and selection frequently determines which mutations will be observed within a malignancy^33^. Genetic “hits” are not limited to somatic mutations and we note that recurrent copy number alterations are more characteristic of invasive breast cancers than recurrent mutations^33^.

We established a case-control study of benign breast biopsy (BBB) samples, The Benign Breast & Cancer Risk (BBCAR) Study. We have performed whole exome sequencing on the benign biopsies of 135 subjects who subsequently developed breast cancer (cases) and from 69 age and race matched controls, who have not developed breast cancer to date. The cases and controls had similar degrees of benign change: non-proliferative or proliferation without atypia. This is an important consideration as these lesions, rather than being obligate precursors, predict risk of the development of breast cancer with an almost equal distribution between the two breasts when a malignancy is diagnosed subsequently^22^. We hypothesized that like the eyelid epidermis, benign breast lesion would also harbor somatic mutations and that by identifying the mutational signatures associated with these mutations we could surmise the mutational processes the tissue had been exposed to. Additionally, we hypothesized that the genetic alterations that differ between cases and controls would provide clues to the molecular alterations responsible for the earliest steps in oncogenesis and would enable the precise prediction of breast cancer risk. To our knowledge, this is the first empirical case-control study that has performed whole exome sequencing (WES) on benign breast biopsies without atypia. In addition to two known mutational signatures, we have identified a novel signature that is significantly (p=0.007) associated with triple negative breast cancer. We have determined that the somatic mutation rate in benign lesions is similar to that of invasive breast cancer. Although the number of cases and controls with copy number variation (CNV) data is limited, we have identified several amplifications and deletions that appear to be exclusive to the cases. We have also determined the mutations that are significantly associated with time to the diagnosis of breast cancer, and mutations common to the benign biopsy tissue and the breast malignancy for the ten cases for which we had matched pairs.

## Results

### Mutation catalogues

Among the 204 samples, 36,801 somatic base substitutions and 2,283 small INDELs were identified. There is a substantial variation in the number of mutations between samples. Cases had a mean of 6.2 mutations/MB (SD 3.6) and controls 6.8 mutations/MB (SD 3). After excluding the synonymous changes, the majority of the remaining mutations were missense mutations (Fig. 1A). No significant difference was observed in the numbers of mutations between the case and control (Fig. 1A). Among the top 20 mutated genes, the case group and control group shared common genes (*MUC17*, *OBSCN*, *FLG2*, *GLTPD2*, *ABCA13, PIK3CA*) (Fig. 1B). Approximately one-fifth of both cases and controls display *PIK3CA* mutations. At a glance, the proportions of nonsense mutations vary between samples too (Fig. 1C). Majority of nonsense mutations were frame shift insertion and stop gain, with some exceptions in a few samples (Fig. S1).

**Fig. 1:**
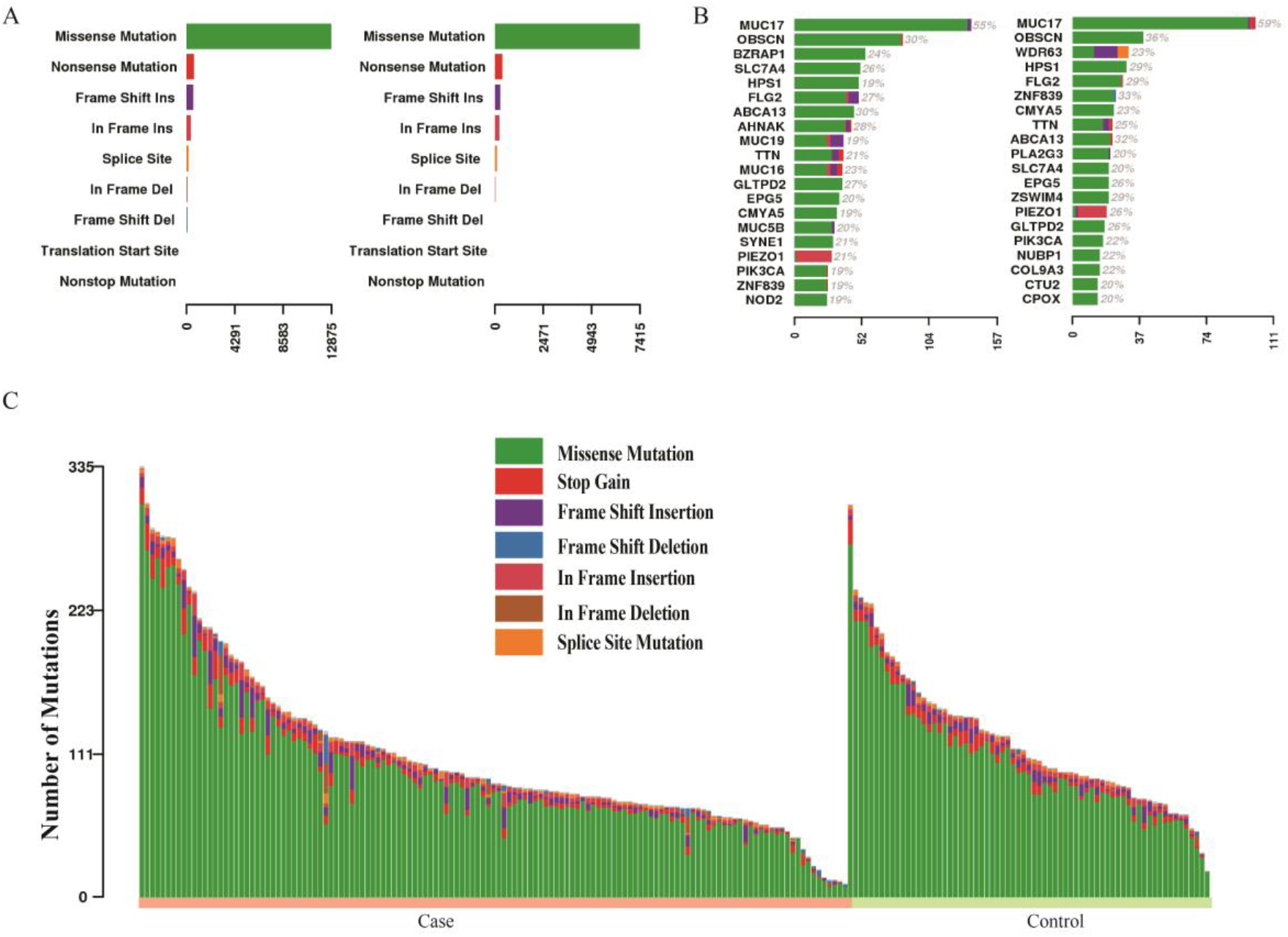
Catalog of somatic mutations in 204 benign breast biopsies. A. Mutation classes in the case group and control group B. The top 20 mutated genes in the case group and control group. C. Catalog of base substitutions, insertions/deletions in the 204 benign breast biopsies. Color codes for mutation class provided in A.

### Genes enriched with mutations in the case

To determine the enrichment of mutations in the case group, a logistic regression model was fit for each gene, with output variable as case/control and input variable as binary variable indicating whether the individual has mutations in the studied gene. The p-values were derived from the fitted models for gene sorting (Fig. 2). Nonsynonymous mutations in four cancer-associated genes, *FLG*, *GNAS*, *CTNNA2*, and *BCORL1*, were more abundant in the case group compared to control. The same analyses including synonymous mutations is presented in Fig. S2.

**Fig. 2:**
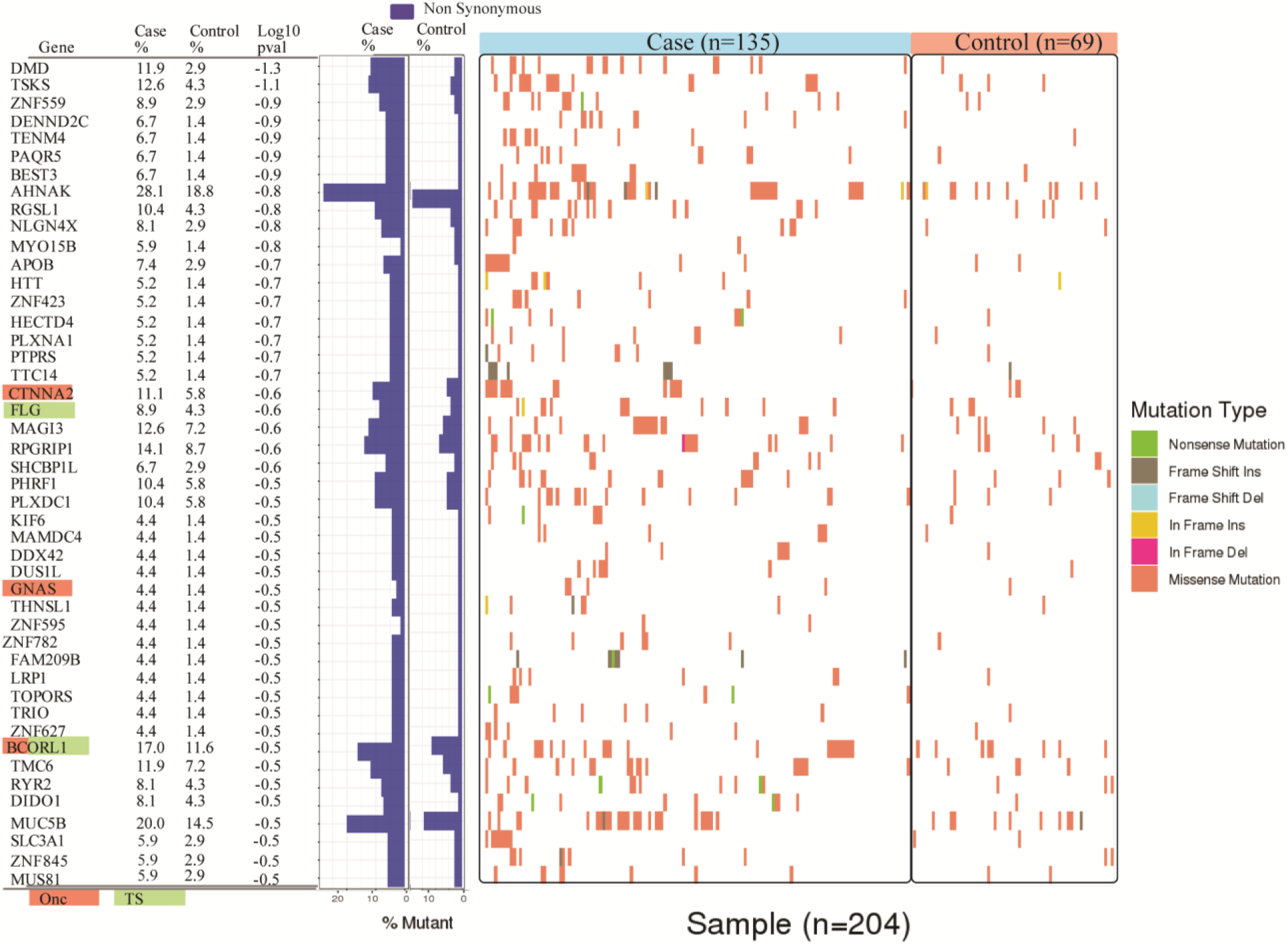
Genetic aberrations that distinguish case and control. For each gene, the percentage of mutated individuals in the case and control are shown. Onc are known oncogenes; TS are the tumor suppressor genes (Cancer Gene Census; https://cancer.sanger.ac.uk/census). The middle panel shows the nonsynonymous rate in each group. In the right panels, each column is imputed with individual’s mutation information, and the color indicates the mutation class.

### Mutational processes

Mutations are non-random and occur within sequence motifs. These motifs provide evidence from which we can infer the process that created the mutations. Recent studies led by investigators at the Welcome Trust Sanger Institute (WTSI) presented the somatic mutation data as a 96-element vector, which captures the immediate 5’ and 3’ neighbors of the mutated nucleotides. Employing non-negative matrix factorization (NMF), 30 “mutational signatures” were produced by these groups^32,34^. Within the BBB cohort, mutational signatures were examined. Three mutational signatures in each case/control group were identified. In both case and control group, we identified the “aging” signature (WTSI Signature 1b; Fig. 3; cosine similarity score: 83.2% for the case and 83.0% for the control), which is the putative result of the hydrolysis 5-methylcytosine. We also identified the “mismatch repair” signature (WTSI Signature 6; Fig. 3; cosine similarity score: 80.5% for the case and 80.1% for the control). A new signature was also identified in each group, both demonstrate an enrichment of C>A mutations with a 5’G or T (Fig. 3).

**Fig. 3:**
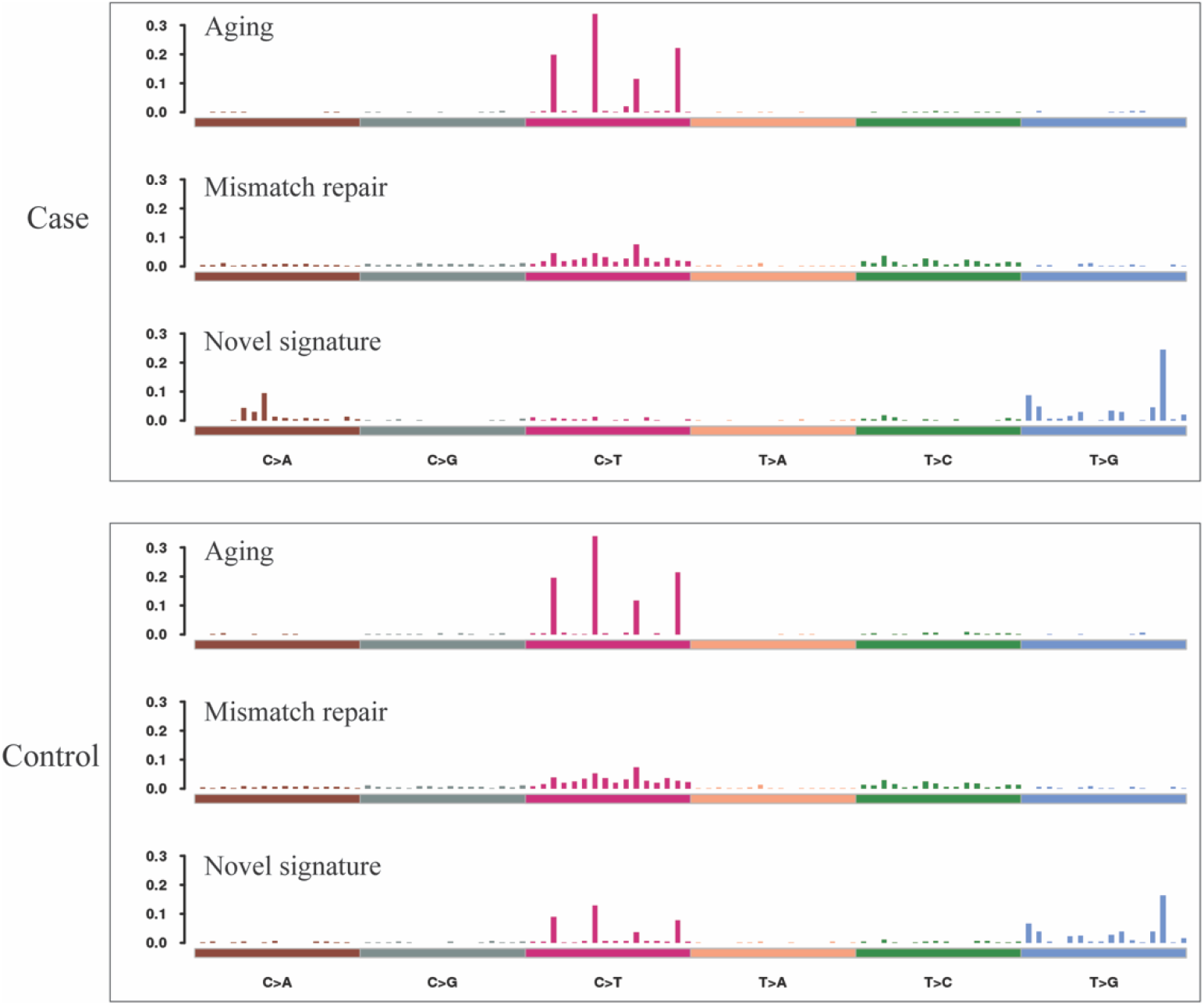
Mutational processes identified in the case group and the control group. The mutations signatures were compared with those of the Welcome Trust Sanger Institute, whose catalog of signature now reaches a total of 40^35^. The aging signature and mismatch repair signature are enriched in both groups. We hypothesize that the novel signature is due to oxidation.

A majority of breast tumors, especially those that are HER2 positive, have been reported to be enriched with mutations hypothesized to result from the action of the APOBEC enzymes ^36^. In our cohort, no tumors were found to be enriched with mutations within in the APOBEC motif nor did we observe either WTSI Signatures 2 or 13 both of which are hypothesized to be the result of the activity of these enzymes. We have also examined the subset of 11 BBB that eventually developed HER2 positive cancer and the subset of 29 BBB that developed cancer within 3 years of biopsy, and we found no APOBEC signatures enriched in these BBB.

The process of deriving mutational signatures is an unsupervised learning process. Pooling the cases and controls together, we derived three signatures in the BBB cohort, namely aging, mismatch repair, and a novel signature. In an association study, we found that the novel signature was significantly (p=0.007) associated with triple negative breast cancer (Fig. 4). In the association study, controlling for the potential covariates of age, menopausal status, and histology class (non-proliferative or proliferative without atypia), the association remains significant (p=0.016), suggesting that novel signature we have identified, is predictive of triple negative breast cancer.

**Fig. 4:**
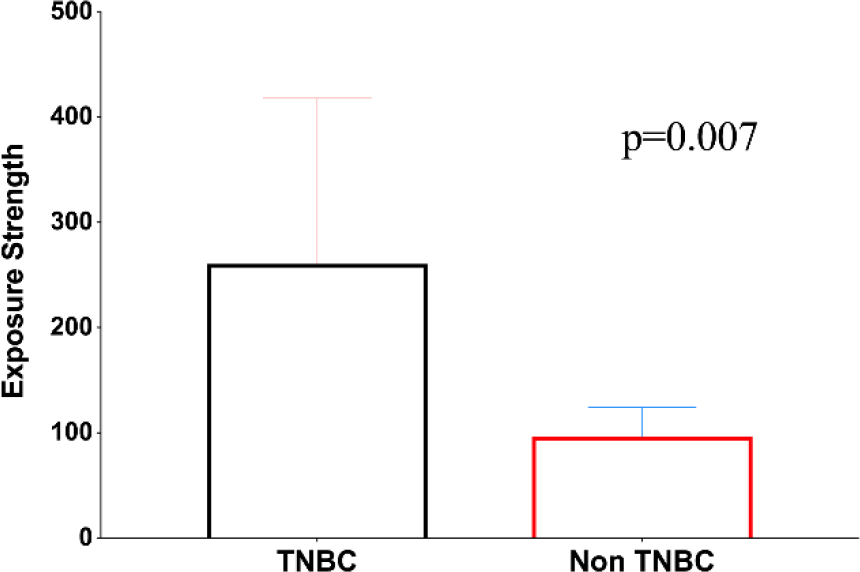
The novel signature is enriched in triple negative breast cancers.

**Fig. 5:**
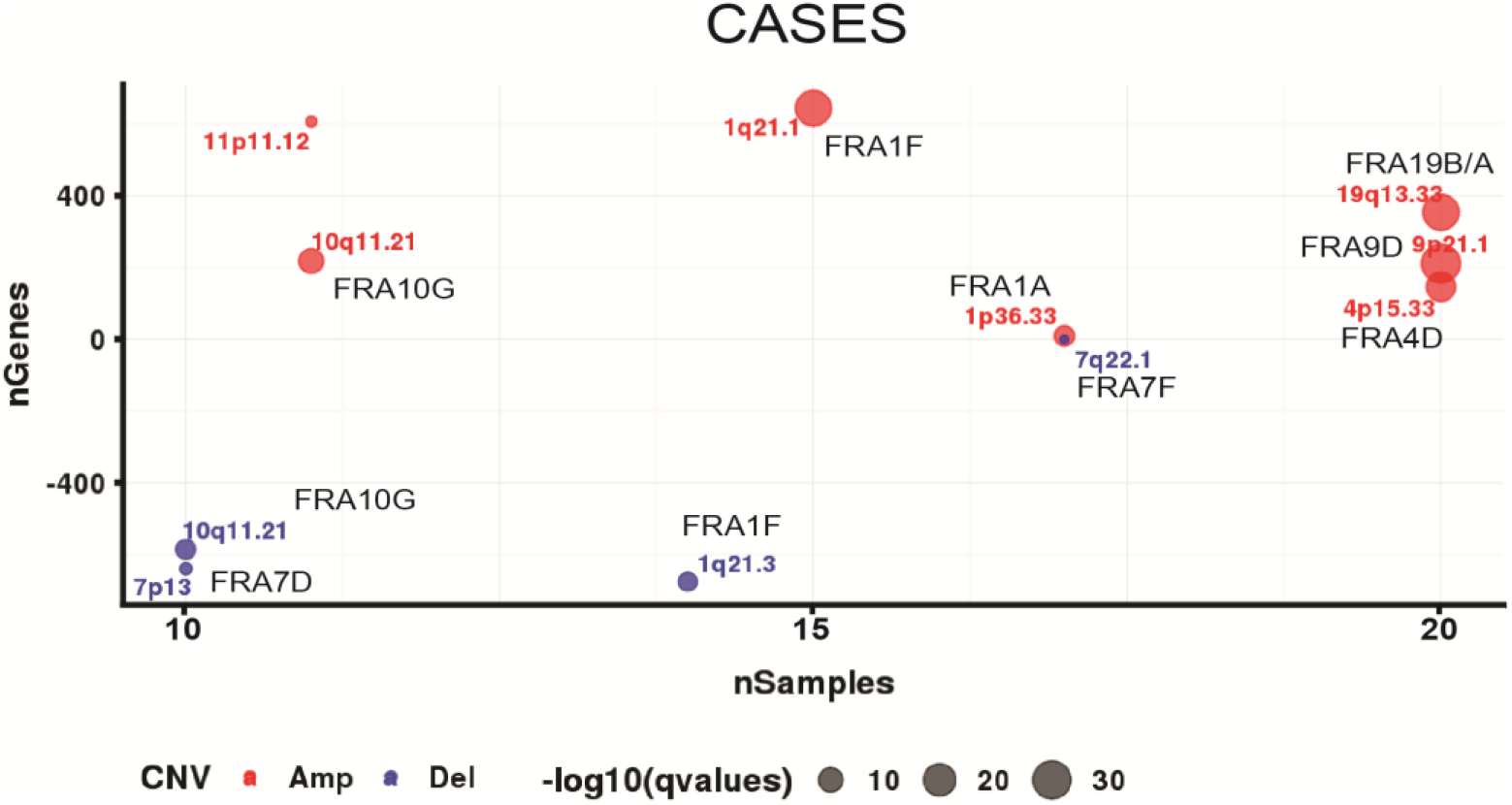
Recurrent somatic copy number variation in the case group. Common fragile sites are indicated in purple

**Fig. 6:**
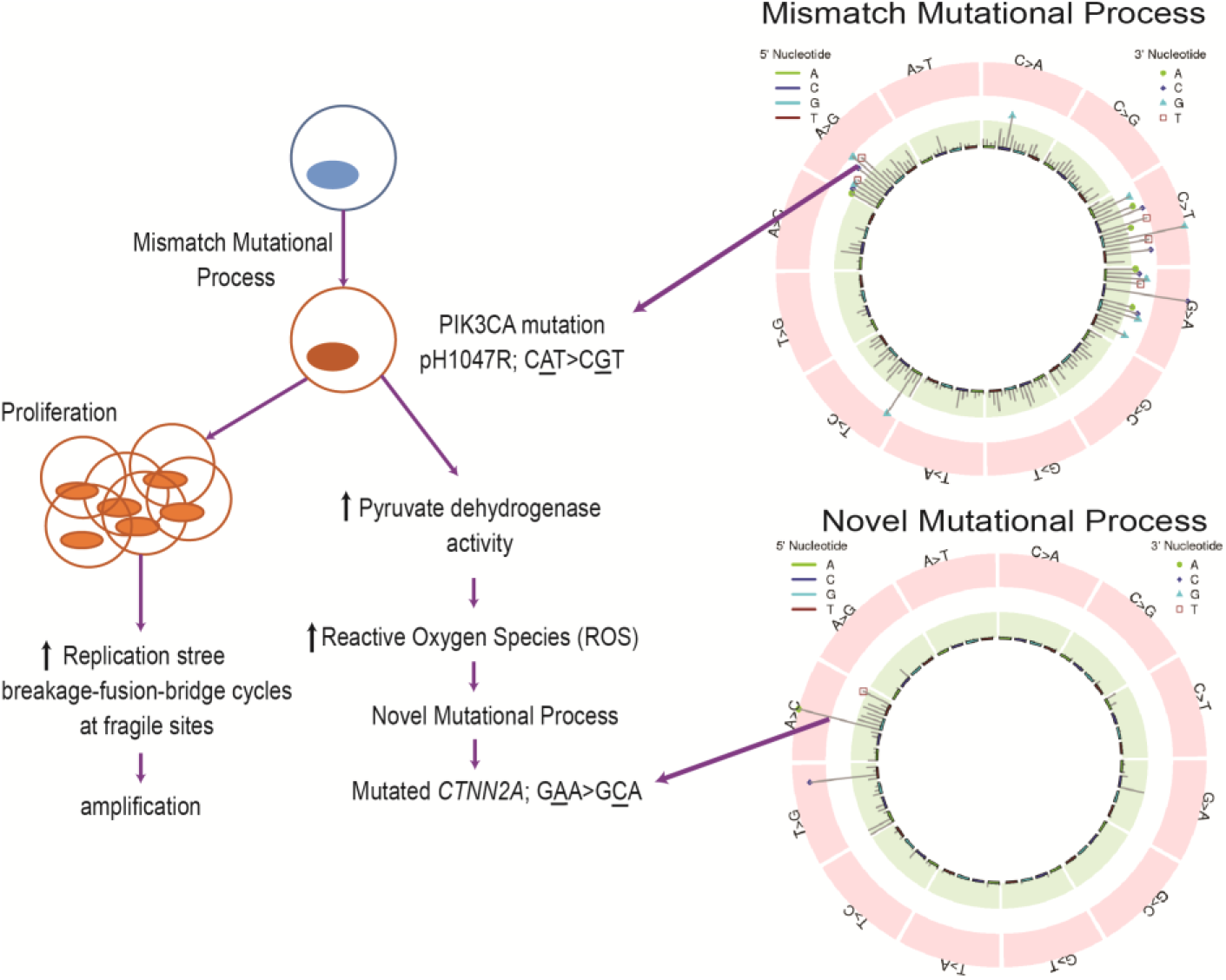
Hypothesis. In illustrations of the mutational signatures/mutational processes, the WTSI 96 trinucleotide format is replaced by a representation of all 192 trinucleotide motifs

### Copy number variation

We employed VarScan2^37^ to study somatic copy number variation in the 26 samples for which we have matched normal DNA. We then applied GISTIC2^38^ to study recurrent CNV. The cytobands we found in the case group, but not in the control group, are presented in **Error! Reference source not found.**. With the exception of 11p11.12, all of the cytobands occur at or immediately next to common fragile sites suggesting that these cells are under considerable replication stress.

### Cancer risk prediction at benign breast biopsy

In an attempt to build a model for cancer prediction at the time of BBB, we fit logistic regression with L1 penalty using the case/control as output variable and the aggregated the number of mutations in each protein domain as a continuous number for input variable. In the evaluation, we performed a bootstrapping by randomly splitting the samples at a 7:3 ratio, and trained the model using 70% of the samples while 30% of the samples were used for prediction. We repeated the process ten times and obtained an AUC of 67% (95%CI: 63.1% to 70.9%) in predicting the cases. Of note, inclusion of clinical characteristics and demographics, including age at time of BBB, age at menarche, age at first live birth, first degree family breast history, histologic variable (proliferative vs non-proliferative), radial scars, and columnar cell alterations (sclerosing adenosis), did not improve the model’s performance.

### Somatic mutations present in both benign biopsy and cancer

We retrieved the paraffin blocks from the cancers that developed in ten of the BBCAR cases. The same procedures as with the BBB were performed, including LCM, DNA extraction, library construction, sequencing, alignment, mutation calling, and variant filtering. Due to the relatively higher allele frequency of mutations in tumor tissues^30^, the read depth cutoff was adjusted to 10, instead of 20^39^. This is a cutoff has been used extensively in other studies. In total, 10,402 mutations were identified in these ten cancer samples. Of these mutations, 957 were observed in both the benign biopsies and cancer tissues. The average allele frequencies for these mutations is 32.2% (SD=18.7%) in the benign biopsy and is 46.7% (SD=17.3%) in the cancer tissues. *FAT1*, *CTNNA2*, and *ATR* among the top ten mutated genes are also are known tumor suppressor genes or oncogenes (Table 1). All six of the *CTNNA2* mutations occur within the motif 5’G<u>A</u>A3’>5’G<u>C</u>A3’. This motif is a predominant feature of our novel mutation signature.

**Table 1:** Somatic mutations present in both benign biopsy and cancer. N = the number of mutations; “Consensus” is the role the mutated gene has been shown to play in oncogenesis per the Cancer Gene Census (https://cancer.sanger.ac.uk/census).

| Gene Symbol | Description | N | Consensus |
| --- | --- | --- | --- |
| *FAT1* | FAT Atypical Cadherin 1 | 8 | TSG |
| *EPG5* | Ectopic P-granules autophagy protein 5 homolog | 7 |  |
| *BZRAP1* | TSPO associated protein 1 | 6 |  |
| *CTNNA2* | Catenin Alpha2 | 6 | OG, TSG |
| *MYH11* | Myosin Heavy chain 11 | 6 |  |
| *ABCA13* | ATP binding cassette subfamily A member 13 | 5 |  |
| *ATR* | ATR Serine/Threonine Kinase | 5 | TSG |
| *DNAH14* | Dynein axonemal heavy chain 14 | 5 |  |
| *ETAA1* | ETAA1 activator of ATR kinase | 5 |  |
| *FBXL18* | F-box and leucine rich repeat protein 18 | 5 |  |

\* TSG =tumor suppressor gene; OG = oncogene

## Discussion

Cancer is thought to occur as a consequence of the progressive accumulation over time of several somatic mutations. Based on epidemiology evidence, Armitage and Doll hypothesized that no more than two genetic events are necessary to create a malignancy, the first event resulting in a faster rate of multiplication that confers a selective advantage to the affected cells^40^. The size of the mutant clone relative to other normal clones then continuously increases and the clinically apparent cancer manifests after the occurrence of a second mutation within the clone. Other evidence suggests that there is a range of “hits” but the number is small with the upper limit probably no more than five^26,27^. Genetic aberrations associated with malignancy occur within normal tissues^30^ and within tissues at the population risk of breast cancer^28^ as well as within lesions at substantial risk^29^. Rohan and colleagues utilized targeted sequence capture and sequence of the DNA from the benign breast disease tissue from a case-control study of women who eventually developed breast cancer compared those who did not^41^. While they identified somatic mutations in a number of genes frequently mutated in breast cancer, no significant differences were identified comparing cases and controls with regard to mutational burden, genes mutated, type of mutation or pathway. No mutations shared between the biopsy and the tumor were observed. We utilized a similar case-control design but employed WES. We rigorously evaluated the sequencing quality, mutation calling, mutation prediction, and concluded that we have obtained a set of valid somatic mutations. We developed a neural network model to predict somatic mutations for the benign biopsies without matched normal DNA, and obtained a precision of 95%. This tool, if validated in an independent collection of tissue, will be of inestimable value as almost every medical facility archives paraffin blocks of benign breast biopsies most of which do not have matched germline DNA. Using the sequencing data produced, we have identified recurrent mutated genes and copy number variations. We also built a predictive model for the risk of breast cancer using the genetic information alone and obtained an AUC of 67%. We have identified a novel signature that is associated with triple negative breast cancer (p=0.007).

### Copy Number Variation/Copy Number Aberration

Single cell sequencing of synchronous DCIS and invasive ductal carcinomas has revealed that copy number aberrations (CNA) are early oncogenic event, i.e., present in *in situ* lesions, and that no additional CNA events are acquired during the transition from in situ to invasive lesion^42^. Recurrent copy number alterations are a characteristic of invasive breast cancers rather than recurrent mutations^43^. CNAs are hypothesized to be the result of short bursts of genomic instability. The mechanisms of punctuated copy number evolution are unknown; telomere crisis has been advanced as a plausible model. Our data clearly show that CNV/CNAs are present in lesions that are markers of risk but not obligate precursors (i.e., non-atypical benign change, with or without proliferation), and that those present in the cases do not overlap with those found in the controls. We also observe that the CNA occur within or adjacent to identified common fragile sites. Common fragile sites (CFS) are regions of recurrent double-strand breaks originally observed in metaphase spreads. The double strand breaks of CFS are associated with replicative stress that can be the result of any of a number of endogenous or exogenous factors that either cause replicative fork collapse or prevent its resolution. Among these factors are depleted deoxyribonucleotide pools, *ATR* dysfunction, activated oncogenes, and inhibition of DNA polymerases. Interestingly, of the mutated genes shared between cases’ BBBs and tumors, two are involved with ATR function: *ATR* itself and *ETAA1* activator of ATR kinase. Both gene products function at stalled replication forks to maintain genomic stability. Amplifications are hypothesized to be the result of breakage-fusion-bridge (BFB) cycles triggered at the induction of fragile sites^44^.

The cytobands at which CNAs are identified in the cases have been associated with invasive breast cancers. One of the amplification outliers identified by Curtis and colleagues using tumors from the METABRIC consortium was within chr19q13.33 and contains 26 genes. No candidate oncogene has yet to be identified within this amplicon, however *TSKS*, a gene more frequently mutated in our cases is among these 26 genes^45^. Chromosome 1q21 is fourth most frequent locus of copy number alterations in cancer^43^. Although no specific oncogene or tumor suppressor gene within this cytoband has been identified yet, a putative oncogene within the chr1q21.1 amplicon is *BCL9*. An analysis of TCGA data by Elarraj and colleagues revealed that 13% of invasive breast cancers harbor an amplification of this gene^46^. These investigators also identified a significant association between *BCL9* gene amplification and mRNA upregulation. This increased expression of BCL9 has been associated with the transition of DCIS to invasive carcinoma, which begs the question as to whether increased expression is also associated with the CNA observed at chr1q21.1 in the BBCAR cases and therefore involved in the transition of normal to at risk tissue.

### Mutational Signatures

We have identified three mutational signatures. Our aging signature is most similar to WTSI Signature 1b, which is associated with aging^25^ and hypothesized to be the result of the hydrolytic deamination of 5-methylcytosine^47^. Our mismatch signature is similar to WTSI Signature 6. *MLH1*-inactivated cancers have a combinations of C>T/G>A and T>C/A>G transitions, which are associated with WTSI Signature 6 and 26, respectively. There are no *MLH1* mutations in either cases or controls. Our novel signature is enriched with T>G/A>C mutations, with 5’G<u>A</u>A3’>5’G<u>C</u>A3’ the most frequently mutated trinucleotide. These transversions are observed *in vitro* when equimolar oxidized dGTP (8-O-dGTP) is included in the nucleotide pool^48^. There is a 4-to 5-fold difference in the mutation rate depending on the sequence context with 5’GAA3’ being a favored context under these experimental conditions. We made an extra effort to look for the mutational signatures associated with APOBEC activity; we were unsuccessful. This may not be so surprising as we are examining relatively early events in oncogenesis. APOBEC mutagenesis (WTSI Signature 2) is a late event in tumor evolution fostering the expansion of subclones^49^. This increasing prevalence is observed for ER-negative breast cancer but not for ER-positive. In addition to being a late effect, a recently published mutational signature study of 1,001 human cancer cell lines and 577 xenograft revealed that APOBEC-mediated mutagenesis is episodic. The initiating factors have not yet been identified but substrate availability and retrotransposon mobilization may contribute^35^. We note that stalled replication forks would theoretically provide single-stranded substrate.

### Somatic Mutations

*PIK3CA* is the most frequently mutated gene in breast cancer; it is observed in 28% of cases (Sanger COSMIC, https://cancer.sanger.ac.uk/cosmic). There are three mutational hotspots, that in the kinase domain of the *PIK3CA* gene, pH1047R, is preferentially selected in invasive breast cancers^33^. MCF-10A-H1047R is a cell line in which there is a knock-in mutation of H1047R in one allele of *PIK3CA*^50^. Comparisons of this cell line to its parent, MCF10A, has revealed the profound effect this single mutation has on the mutation of other genes, gene expression, protein expression, AKT and MAPK pathway activation and metabolism^51^. Whether this mutation can single-handedly transform cells is debatable. The lineage tracing studies of Koren and colleges demonstrated that expression of PIK3CA H1047R resulted in tumor formation in both luminal and basal lineages, as well producing multipotent stem-like cells^52^. However, earlier studies carried out by Gustin and colleagues using the knockin cells demonstrated some features of oncogenic transformation but these investigators argued that when the when the mutant gene is under the control of its native promoter the mutation is not tumorigenic^50^. *PIK3CA pH1047R* mutation is observed in one-fifth of the BBCAR cases and controls that we have sequenced suggesting it is an early event and may play a role in the genesis of the benign lesion. All of the mutations are A to G in the motif: 5’CAT3’>5’CGT3’. This mutation motif occurs only in our mutational signature, which is most similar to WTSI Signature 6^53^. We hypothesize that this mutation may represent the first hit of Armitage and Doll’s 1957 hypothesis^40^. As this *PIK3CA pH1047R* is present in both cases and controls, the additional hits that lead to malignancy must be exclusive or overrepresented in the cases.

The MCF-10A-H1047R cell line was initially thought to be isogenic with its parent, MCF 10A. However, sequencing of the two cell lines revealed 43 nucleotides are mutated between the within a few passage in cell culture. This suggests that the H1047R mutation may induce genomic instability. The 48 mutated nucleotides appeared in 58 genes; 13 of these genes were also mutated in our gene sets.

The gene at the top of the list of differentially non-synonymously mutated genes is *DMD* (p=0.1). Mutations within this gene have been associated with the development of Dystrophinopathies^54^, rather than with that of malignant tumors. However, there is emerging evidence to suggest that DMD may function as a tumor suppressor^55^. A recently published pan-cancer molecular study of gynecologic and breast cancers revealed that one of the genes displaying significant copy number loss when comparing pan-gynecologic to non-gynecologic is *DMD*^56^. Another study using data from cBioPortal, which collates next generation sequencing data from The Cancer Genome Atlas (TCGA) and the International Cancer Genome Consortium (ICGC), discovered a median mutation frequency of almost 4% in *DMD* in sporadic breast cancers^54^. This frequency is identical to that of more well know tumor suppressor genes in breast cancer, *PTEN* and *ARID1A*. In induced pluripotent stem cells, loss of dystrophin leads to genomic instability^57^.

### Somatic mutations and time to breast cancer development

We observed a statistically significant association between time to cancer development and *INO80D* mutations. INO80D is one of 13 subunits of the ATP-dependent chromatin remodeling complex INO80^58^. INO80 exhibits a plethora of functions including transcriptional regulation, DNA replication and repair ^59^. Mammalian INO80 stabilizes stalled replication forks and enables their restart following release from replication arrest and therefore, not surprisingly, INO80-deficient cells evidence decreased fork stability, fork collapse and formation of double-strand breaks in response to replication stress^58^. The observation that *INO80D* mutations are statistically significantly associated with time to breast cancer development suggests a possible tumor suppressor role. While no information is available regarding the D subunit, *INO80C* has been identified as a novel tumor suppressor in *KRAS* mutant colorectal and pancreatic tumor xenografts^60^.

We also observed a statistically significant association between time to cancer development and *TENM4* mutations. TENM4 is one of four highly related human teneurin paralogues^61^. The teneurins are large transmembrane glycoproteins with many structural similarities to Notch which include their analogous transmembrane localization, their capability to dimerize upon ligand interaction, the presence of multiple epidermal growth factor-like (EGF) repeats in their extracellular domains, and their processing into multiple domains through proteolytic cleavage^62^. A catalog of chromosomal translocations involving the teneruins discovered that 18/32 (56%) of these occurred in breast cancer (16 of which involved *TENM4*), and most were intrachromosomal (22/32, 69%)^62^. However, which, if any, of these has functional consequences remains a matter of conjecture.

### Gene mutations shared between the cases BBB and tumors

The ten most frequently mutated genes shared between the cases’ BBB and tumors are given in Table 1. *CTNNA2*, *ATR* and *ETAA1* are discussed below. *FAT1* has the most mutations, which is interesting as this same gene was shown to have a statistically significant excess of inactivating mutations across all classes in the sun-exposed, physiologically normal epidermis study^30^. *FAT1* encodes a cadherin-like protein and its inactivation via mutation may lead to tumorigenesis by multiple avenues^63,64^. It is mutated in approximately 2.5% of invasive breast cancers NOS but 11% of metaplastic breast cancers. It is located at chromosome 4q35.2 just beyond the breast cancer deletion outlier at 4q35.1 identified by the METABRIC consortium investigators^65^.

### Hypothesis

We have utilized the above data to develop an hypothesis of the early oncogenic steps for a subset of the cases. *PIK3CA* pH1047R mutations are created by the mutational process which, by it is similarity to WTSI Signature 6, is associated with Mismatch Repair deficiency^53^. Extant nucleotide pools are inadequate to supply the rapid proliferation engendered by the *PIK3CA* mutation resulting in replication stress. This leads to breakage-fusion-bridge cycles at fragile sites^66,67^ and the amplifications we have observed in the CNA data in the cases. *PIK3CA pH1047R* also results in metabolic reprograming with upregulation of pyruvate dehydrogenase (PDHc), which produces mitochondrial acetyl-CoA^68^. PDHc also produces superoxide/H_2_O_2_^69^, which, via the Fenton Reaction, is capable of oxidizing dGTP within the nucleotide pool^70^. 8-O-dGTP is incorporated across from dA during replication and when not repaired leads to the A to C mutations in the motifs^48^ we observe in our novel mutational signature. *CTNNA2* mutations occur in the cases with twice the frequency of the controls (Table 2). The overwhelming majority of mutations occurs within one of the most frequent motifs of our novel mutational signature: 5’G<u>A</u>A3’>5’G<u>C</u>A3’. It is mutated in 6 of the 10 benign-malignant pairs and mutations only occur in the 5’GAA3’>5’GCA3’ motif in these pairs. *CTNNA2* is a tumor suppressor gene and its mutated form has been identified as a putative driver of subclonal expansion^49^. Replication stress leads to stalled replication forks and dysfunction of *ATR* or *ETAA1*, two of the mutated genes shared by cases’ BBB and tumor, if unable to stabilize the forks and allow time for repair would lead to further genomic instability^71^. *ATR* also specially regulates fragile site stability^72^. This is a rudimentary sketch, which will certainly be edited as more information becomes available, and it only represents a minority of cases. Nevertheless, many of it features can be tested.

## Materials and Methods

### At the Northwestern Feinberg School of Medicine, we designed a case-control study of benign breast biopsy (BBB) samples (Benign Breast & Cancer Risk, BBCAR Study)^73^

We have retrieved the BBB paraffin blocks of subjects who subsequently developed breast cancer (cases) and from age matched controls, who have not developed breast cancer to date. The participants have provided consent for use of their benign and cancer samples, have completed a risk questionnaire, and are contacted periodically to confirm that controls have not transitioned to cases. A subset of 135 cases, matched to 69 controls were selected for whole exome sequencing (WES) (Supplementary Materials and Methods). The cases and controls were also matched by categories of benign change: non-proliferative or proliferation without atypia. DNA was isolated from the laser captured, microdissected (LCM) epithelium and sequenced using the Illumina HiSeq4000. Whole exome sequencing shave been conducted with sequencing depth of 80-100× and 80-90 million sequencing reads were generated for each sample (Supplementary Materials and Methods).

### Our initial objective was to develop and test a predictive model for somatic mutation identification

One of the most significant challenges for studies seeking to identify somatic mutations in archival tissue samples is that matched germline DNA is unlikely to have been collected and stored. Therefore, to prepare for a ground truth, previously consented donors were re-contacted and saliva specimens were requested for normal DNA sequencing. To date, matched germline DNA has been obtained for 26 (12.6%) of the 204 BBB specimens; WES has been performed on these specimens as well. We systematically evaluated multiple machine learning models and adopted multiple perceptron layer for somatic mutation prediction. Features in the prediction model included intrinsic sequencing features, such as mutation allele frequency, depth of reference reads, number of appearances in the cohort as well as published collated data providing the frequency of the variant in the population and predictions of the impact of amino acid changes on the structure and function of the encoded protein. The model obtained an accuracy of 95% for somatic mutation in the test set (Supplementary Materials and Methods).

### Orthogonal SNP array genotyping was performed to compare and validate the performance of mutation calling and mutation prediction

Technical validation was performed for 17 of the 26 specimens for which matched germline data was available, and 3 of the specimens without matched germline, using the Infinium Exome-24 v1.1 beadchip (Supplementary Materials and Methods). Multiple somatic mutation callers were also carefully evaluated and Mutect2 was selected as our final mutation caller (Supplementary Materials and Methods). Overall, we achieved accuracy of 85.42% in identifying somatic mutations for the samples with matched germline DNA available. Compared to this performance, the ones without germline DNA available, has an accuracy decrease of 5.42% in terms of identifying true somatic mutations. To study the transitions from benign biopsy to cancer, we further sequenced 10 cancer samples that matched to the cases.

### Using both aligned reads and identified mutations, we studied the genetic aberrations that distinguish cases from controls, including mutations and copy number variation (CNV)

We identified the somatic mutations or CNV that were common to both the cases’ benign biopsy tissue as well as their malignant lesions for the ten cases in which we had both tissues available. P values relating to the case or control group were calculated with the use of Chi-square test or logistic regression. We also studied the mutations to enable the discovery of mutational signatures. Lastly, we evaluated machine learning models and features for breast cancer risk prediction for the cohort. Benjamini-Hochberg method was applied to convert the two-sided P-values to False Discover Rate (FDR) for multi-comparison correction.

## Author contributions

ZZ, SAK and SEC conceived the study. AS performed laser capture microdissecion and extracted the DNA. PS administered the questionnaires, organized the clinical data, and contacted the subjects for saliva donation and cancer status confirmation. XX performed the sequencing. ZZ and AV carried out the sequence alignment, quality assessment, and mutation calling. SEC and ZZ wrote the manuscript. ZZ and XL performed the statistical analysis. YL reviewed all analyzed data. SAK was responsible for the clinical study. All authors discussed the results, revised and approved the manuscript.

## Supplementary Materials

**Fig. S1:**
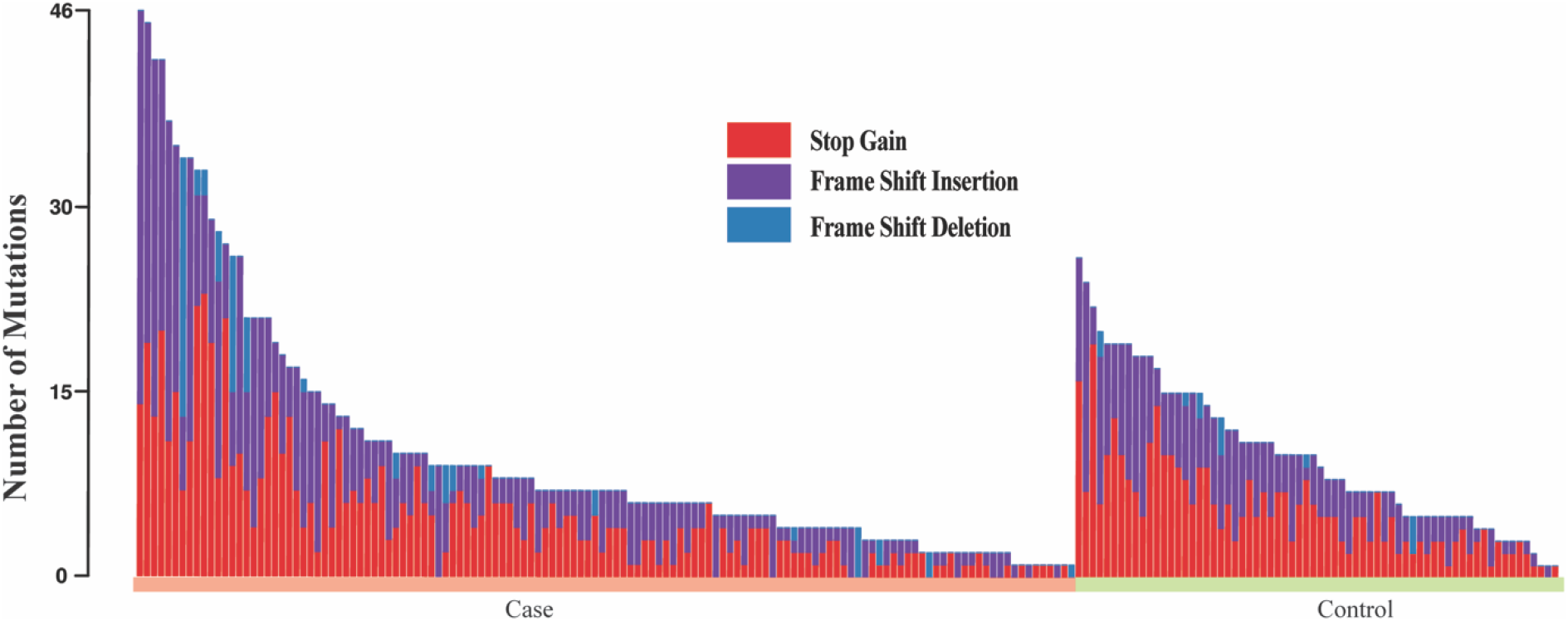
Catalog of nonsense mutations in the 204 benign breast biopsies

**Fig. S2:**
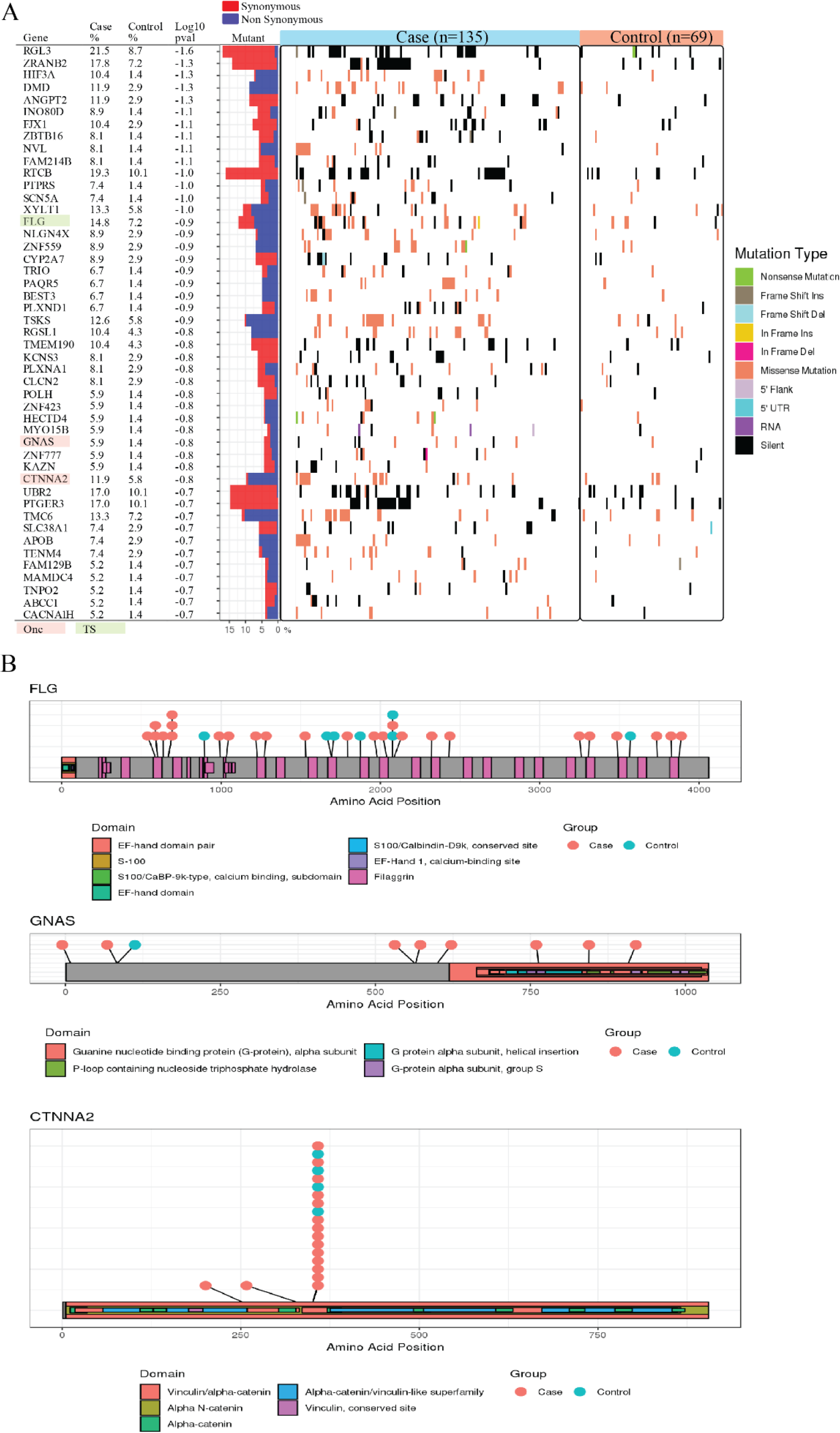
Genetic aberrations that distinguish case and control and mutational positions. A. For each gene, the percentage of mutated individuals in the case and control were shown. The Onc is the known oncogenes; the TS are the tumor suppressor genes. The p-values were derived using the case/control as output and the mutated individual as inputs in logistic regression. The middle panel shows the synonymous versus nonsynonymous rate. In the right panel, each column is an individual, and the color represents the mutation class. B. Position of mutational alterations in the protein structure of FLG, GNAS, and CTNNA2.

### Study design

The Enterprise Data Warehouse (EDW) is a joint initiative across the Northwestern University Feinberg School of Medicine and Northwestern Memorial HealthCare^74^. Using the database, we designed a case-control study to test the hypothesis that mutations present in the benign breast biopsies of women, who went on to develop breast cancer years later, are different from those who did not develop this disease, and these mutations could be used as markers of the risk of development of breast cancer (Fig. S3A). The assumption underpinning the proposed research is that the breast tissue of the cases harbors genetic alterations predictive of the eventual development of breast cancer that were present years before the malignancy could be detected. Additionally, it is assumed that although a specific geographic area of the breast, i.e., the breast tissue in the archival block, is available for assessment, any genetic alterations identified that were predictive would predict generalizable, that is, bilateral risk. Lastly, it is assumed that the genetic alterations are not simply those that predisposed or led to the benign lesion. The second of these assumptions presumes a field cancerization that encompasses both breasts and that predates the detection of the malignancy.

**Fig. S3:**
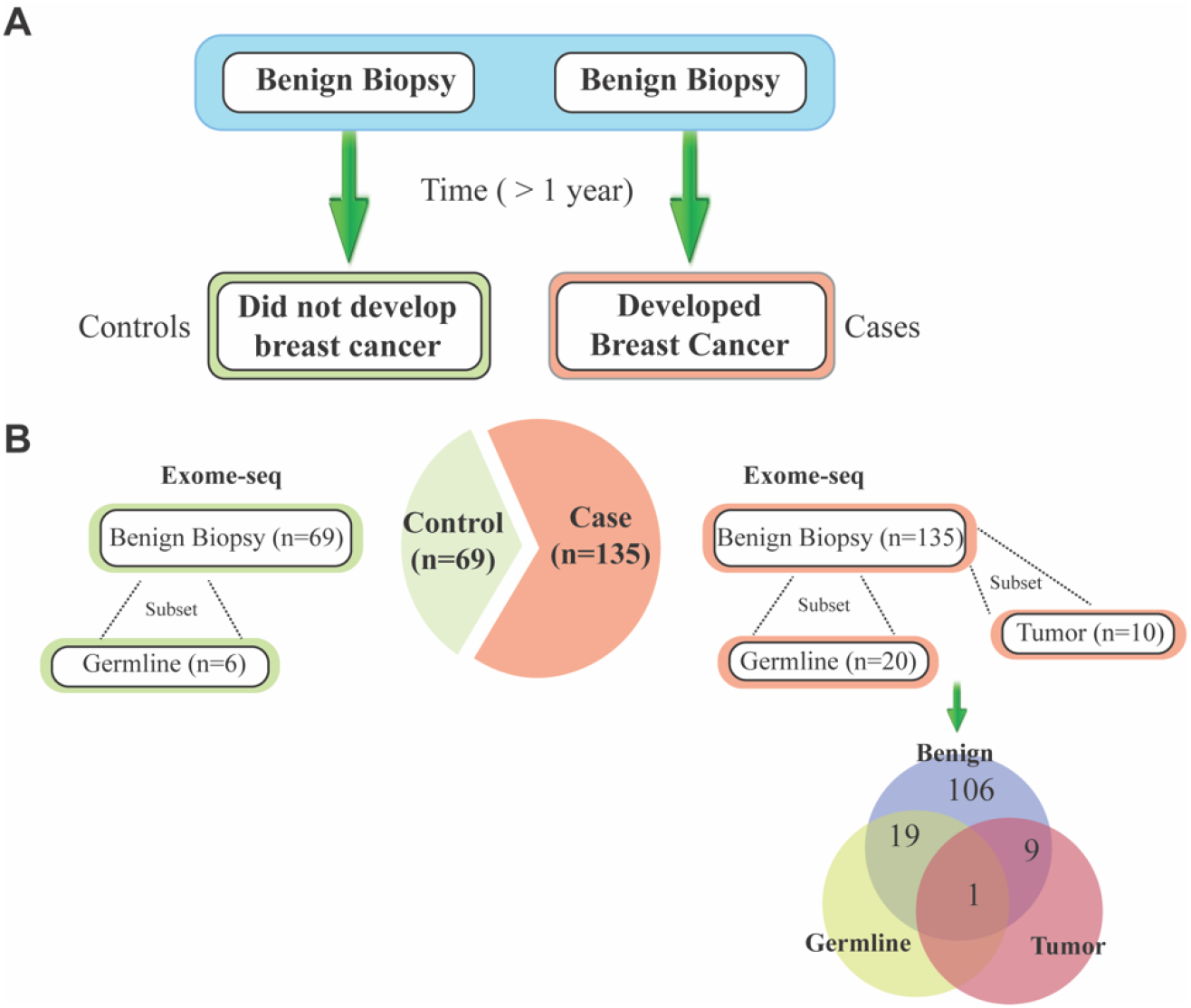
Study design of the study A. Design of the BBCAR study. B. Study Design illustrating the number of sequenced samples.

We identified subjects for a case-control study of benign breast biopsy tissues. Cases (n=135) are women who have undergone a breast biopsy with a benign result on histology that predates the diagnosis of breast cancer by at least one year. The median interval from benign biopsy to the diagnosis of cancer is 7.3 (SD=4.4) years. Controls (n=69) are women who have not developed breast cancer matched for age (± 2 years), duration of follow-up, race, and histology (Table S1). Controls were verified for no cancer development at 08/14/2018. To rule out the germline variants, 16 and 10 matched saliva were collected for germline DNA extraction in the case and control group, respectively (Fig. S3B).

**Table S1:** Distributions of demographic data and tumor characteristics between the Case group and the Control group. Student’s t-tests were performed for continuous variables and Pearson’s Chi-squared tests were performed for categorical variables.

|  |  | **Case (135)** | **Control (69)** | **P-value** |
| --- | --- | --- | --- | --- |
| **Age (SD)** |  | 49.7 (9.9) | 49.8 (9.6) | 0.96 |
| **Menopausal status N (%)** |  |  |  | 0.87 |
| Pre | 114 | 76 (56.3%) | 38 (55.1%) |  |
| Post | 90 | 59 (43.7%) | 31 (44.9%) |  |
| **Class N (%)** |  |  |  | 0.78 |
| *Class 1 | 115 | 79 (58.5%) | 40 (58.0%) |  |
| *Class 2 | 75 | 51 (37.8%) | 25 (36.2%) |  |
| NA | 10 | 5 (3.7%) | 4 (5.8%) |  |
| **ER status N (%)** |  |  |  |  |
| Positive |  | 109 (80.8%) |  |  |
| Negative |  | 23 (17.0%) |  |  |
| Low |  | 3 (2.2%) |  |  |
| **Follow-up years (SD)** |  | 7.31 (4.4) | 16.6 (5.4) | <0.01 |
| **Has matched germline** |  | 20 (14.8%) | 6 (8.7%) |  |
| **Has matched cancer (%)** |  | 10 (7.4%) |  |  |

\*class 1/non-proliferative: "Non-proliferation" and "Benign, NOS"
\*class 2/proliferative: "Ductal Proliferation without atypia"

### Sample collection

Benign biopsies retrieved from the consented patients have been reviewed and confirmed by the collaborating pathologist. A trained interviewer contacted, consented, and completed a detailed questionnaire with the patient. Biopsy sections were reviewed by the pathologist for adequacy and classified as “non-proliferative” (class1), “proliferative” (class 2), or “atypical proliferation”. The best matched areas of non-proliferative and typical proliferative changes for the case-control pair were used for this study. Specimen retrieval, storage and processing procedures were carefully designed with attention to blinding the investigators. Ten 10-micron sections per sample were cut from formalin-fixed, paraffin-embedded (FFPE) tissue blocks, and the matched areas of interest isolated by laser capture microdissection (LCM). In detail, we took slides to Center for Advanced Microscopy (CAM) and used Ziess palm microscope to micro-dissect and collect areas of interest in 500μl adhesive cap (AdhesiceCap 500 opaque-Zeiss order number 415190-9201-000).

### Library construction and sequencing

Total genomic DNA have been extracted from the LCM samples. Tissue DNAs were extracted by Qiagen AllPrep DNA/RNA FFPE it (Cat. No. 80234). DNA concentration was measured by nanodrop. Greater than 300ng of DNA has been isolated from 89% of the specimens processed to date. The concentration and quality of gDNA samples was first assessed using Agilent 4200 TapeStation. Then 100-200 nanograms of DNA per sample were used to prepare single-indexed cDNA library using SureSelectXT Human All Exon V6 (58Mb) (Agilent). The resulting libraries were assessed for its quantity and size distribution using Qubit and Agilent 2100 Bioanalyzer. Two hundred pico molar per liter pooled libraries were utilized per flow cell for clustering amplification on cBot using HiSeq 3000/4000 PE Cluster Kit and sequenced with 2×75bp paired-end configuration on HiSeq4000 (Illumina) using HiSeq 3000/4000 PE SBS Kit. A Phred quality score (Q score) was used to measure the quality of sequencing. More than 90% of the sequencing reads reached Q30 (99.9% base call accuracy). With the goad of sequencing at 100X, the final average sequencing depth was 69X, and there were 80-90 million sequencing reads per sample. To note, after the first round of sequencing, 27 samples had the coverage under 50X, and were re-sequenced for deeper coverage under the same protocol.

### Parallel alignment of whole exome analysis

We adapted the most commonly used open source software for genome alignment and variant calling. Read alignment and variant calling were performed according to the Broad Institute’s Genome Analysis Toolkit (GATK) best practices pipeline^75^. Reads were aligned to the human reference genome (hg19) using Burrows-Wheeler alignment^76^ and Picard 2.6 was subsequently used to sort reads and mark duplicates (Fig. S4). To reduce systematic errors, sorted BAM files were separately generated based on the sequence lane that the reads were generated. By doing so, various technical features that are associated with lane-specific artifacts can be filtered during duplicate marking and base recalibration steps. Base recalibration was done using the GATK 3.6 using dbSNP build 138 as a training set. Mutations were called and filtered using MuTect2 in the GATK package. To capture recurrent technical artifacts, we generated a Panel of Normals (PON) for Mutect2 analysis using the sequenced 26 germline DNA. The PON is created by running the variant caller Mutect2 individually on the normal samples and combining the resulting variant calls with the criteria of excluding any sites that are not present in at least 2 normals. To obtain a set of mutations with highest sensitivity, VarScan2^37^ and VarDict^77^ were also applied for mutation calling. To further ensure a high precision call rate, we filtered all mutations with read depth less than 20. After filtering, mutations were then annotated using SNPEFF^78^, VEP ^79^, and ANNOVAR^80^.

**Fig. S4:**
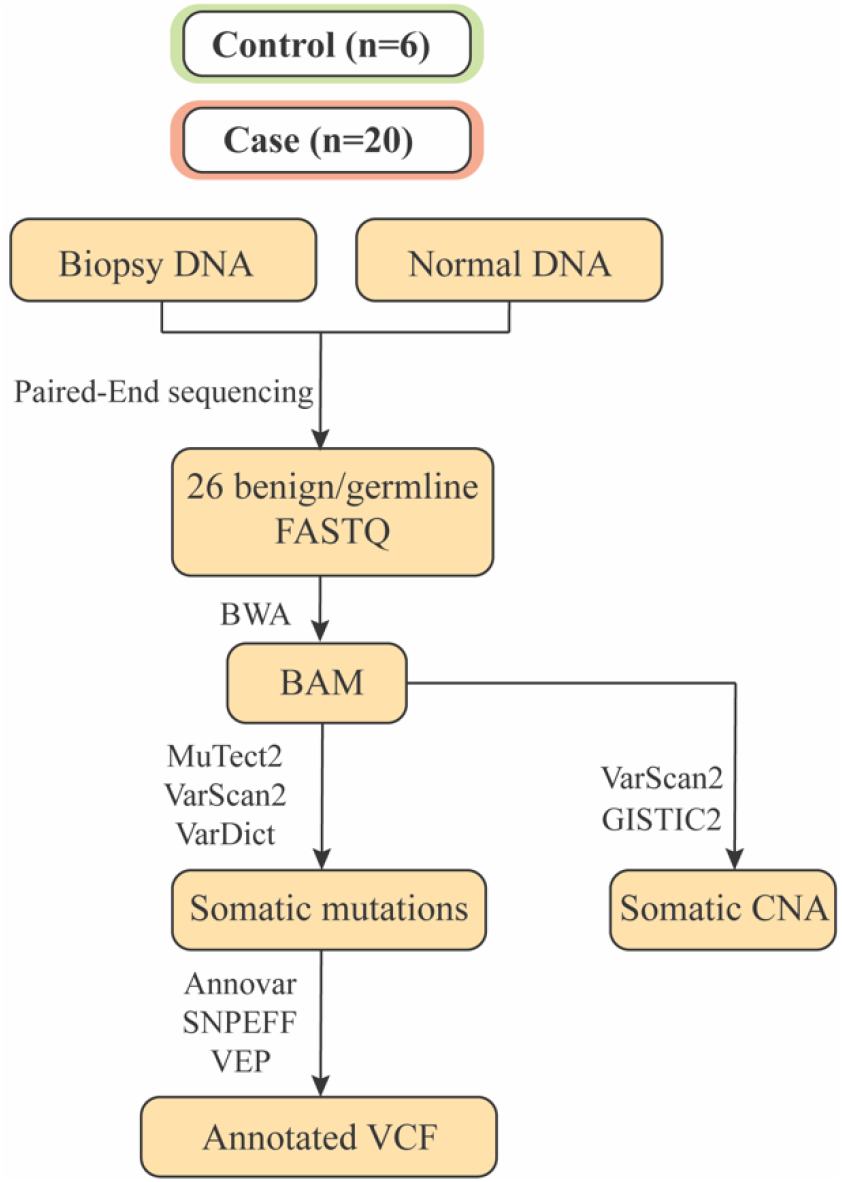
Workflow to call somatic mutations and somatic CNA.

### Microarray genotyping

To evaluate the performance of called somatic mutations, a subset of samples was separately LCM dissected, and the extracted DNA were genotyped using Infinium Exome-24 Kit, which covers 240,000 markers in a catalog of exome variants. To note, the quality control sample had a call rate of 99.34%. During the QC period, no samples failed restoration. For quality control, three samples were repeatedly genotyped twice, and R-square rates were calculated for the overlap. The reported R-square rates are 98.86%, 99.29%, and 99.39%. The high overlap rates indicate a high stability of calling variants from our DNA. Genotyped variants from the 17 samples were mapped to genomic assemblies (hg19). To note, only the calls with GCScore larger than 0.15 were retained. The coordinates that appear both in genotype array and somatic mutations called by MuTect2, VarScan2, or VarDict were retrieved. The allele frequencies derived from both technologies were compared. The overlap number between array and MuTec2, VarScan2, VarDict are 384, 963, and 124511 respectively. When the allele frequency difference is larger than a certain threshold, we consider the call of somatic mutation as a false positive. With different allele frequency cutoffs, we plotted the accuracy rate in Fig. S5. In different thresholds, Mutect2 consistently has a better performance than VarScan2 and VarDict. When the cutoff is 25% (half of 50%), in other words, mutation calls with allele frequency difference larger than 20% between the two technologies were considered as wrong calls, the accuracy of using Mutect2 was 85.42%. Based on the consistent high accuracy rate, we decided to use MuTect2 for further studies.

**Fig. S5:**
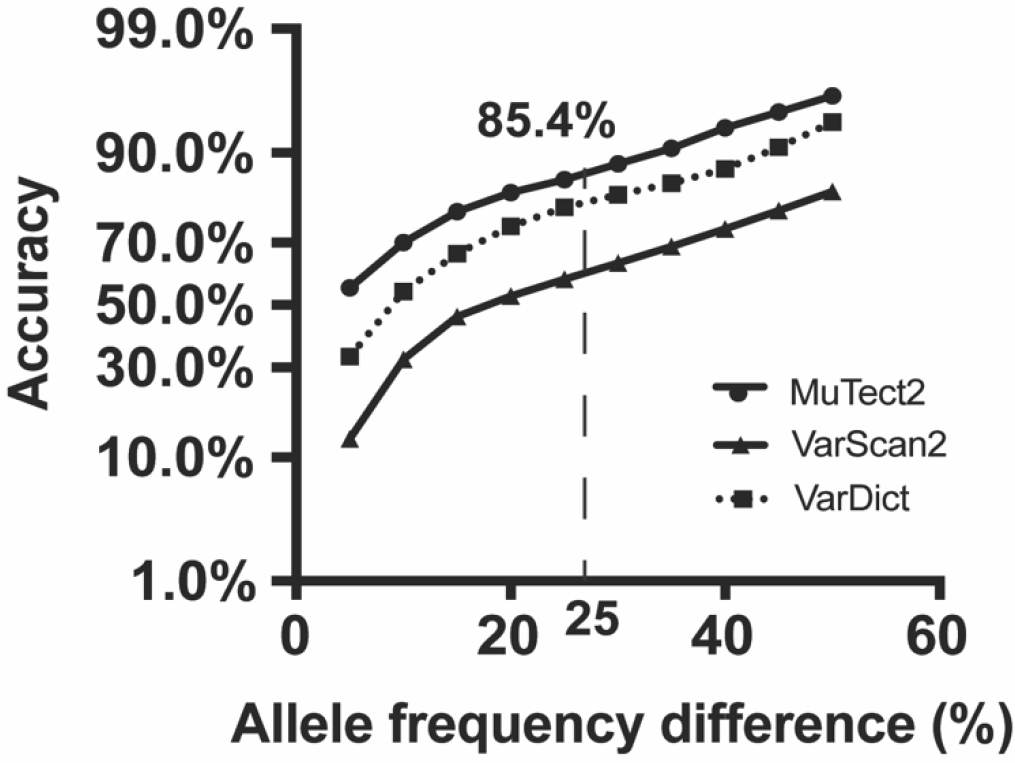
Comparison of the called mutations between genotype array and WES. Somatic mutations were called using Mutect2, VarScan2, and VarDict. With different allele frequency difference cutoff, we obtained different accuracy. The accuracies were plotted.

### Predictive model for somatic mutations

Our initial objective was to develop and test a predictive model for somatic mutation identification. MuTect2 is known as one of the most reliable and sensitive cancer somatic mutation callers^81^. In previous section, we have learned that mutations identified by MuTect2 have a higher accuracy than the other callers. In this study, MuTect2 was used to call somatic mutations from the 26 benign biopsies and matched normal germline DNA. To reduce the false positive call rates, mutations that appear in dbSNP with ANNOVAR^80^ index files (after removing those SNPs < 1% minor allele frequency (or unknown), or mapping only once to reference assembly, or flagged in dbSnp as "clinically associated") and not in COSMIC database (version 80) were labeled as germline variants. The called mutations from these 26 matched samples were used as gold standard for the predictive model training and testing. The called somatic mutations were randomly split to cross validation set and holdout test set based on a 7:3 ratio.

Tools predicting somatic mutations in matched normal free samples have been developed^39,82^. However, the developed tools attempted to predict somatic mutations in tumor-only samples. Tools that have been developed and validated using the mutations derived from tumor samples cannot be applied to benign biopsies directly, mostly due to the different feature landscapes in mutations derived from tumors and biopsies. For example, allele frequency derived in tumor samples are expected to be higher than allele frequencies in benign biopsies^30^. In order to predict somatic mutations in the benign-only biopsies without matched normal DNA, in this study, we attempt to develop and evaluate a new predictive model to predict somatic mutations in benign biopsies.

In total, 31 features (Table S2) were retrieved or developed to create the predictive model for somatic mutation identification. Multiple tools have been developed for potential pathogenicity prediction. These tools consider either the protein structure, population frequency, or evolutionary factors ^83^. Varies functional annotation or toxicity scores were derived from ANNOVAR^80^, COSMIC (https://cancer.sanger.ac.uk/cosmic), dbSNP/common (https://www.ncbi.nlm.nih.gov), along with intrinsic sequencing features, such as mutation allele frequency, depth of reference reads, number of appearance in the cohort. Not all mutations were annotated in each of the database, However, missing data that appear in more than one feature could bring challenge to some of the classifiers (e.g. logistic regression). Considering that the features are a mix of continuous number, binary feature, and categorical variables, Multivariate Imputation by Chained Equations (MICE)^84^ was used to impute the missing values. In detail, 20 sets of data were imputed with the iteration equal to 20. Consolidating the 20 sets of data into one, mean number was calculated for the continuous variables and mode was calculated for the categorical variables.

**Table S2:**
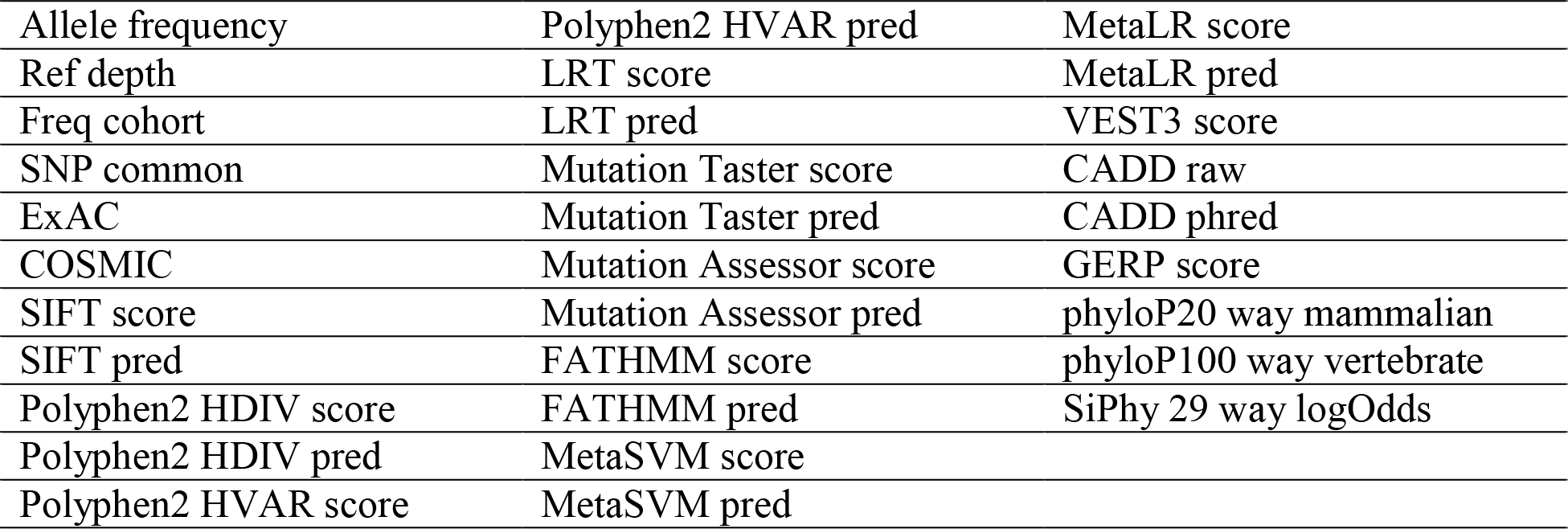
List of features used in the predictive model for somatic mutation identification. ‘Allele frequency’ is the mutation allele frequency, ‘Ref depth’ is the read depth of the reference allele, ‘Fre cohort’ is the number of appearances of this mutation in the cohort. SNP common is a binary variable indicating that the mutation appears in the database of dbSNP after removing those flagged SNPs (SNPs < 1% minor allele frequency (MAF) (or unknown), mapping only once to reference assembly, flagged in dbSNP as “clinically associated”). ‘COSMIC’ is a binary variable indicating whether the mutation appears in the COSMIC version 80. The other features were derived from functional annotations from functional annotation tools.

Utilizing the derived features, we evaluated multiple linear and nonlinear machine learning models for somatic mutation prediction. The germline variants were treated as predictive negative, while the somatic mutations were treated as predictive positive. Grid search was applied to tune each model’s parameters within the five-fold cross validation set. To reduce the risk of having false positives, precision was used as selection criteria for parameter tuning and model selection. Once the model was tuned, it was applied on the holdout test for precision. Recall, F-measure, and AUC score reporting. The model with highest precision was selected as our somatic mutation predictive model. To maximize the prediction power, we evaluated various machine learning methods, including penalized logistic regression (LR), linear SVM, random forest classifier (RFC), gradient boosted tree (GBT), k-nearest neighbor algorithm (K-NN), SVM with rbf kernel, and multiple layer perceptron (MLP).

**Fig. S6:**
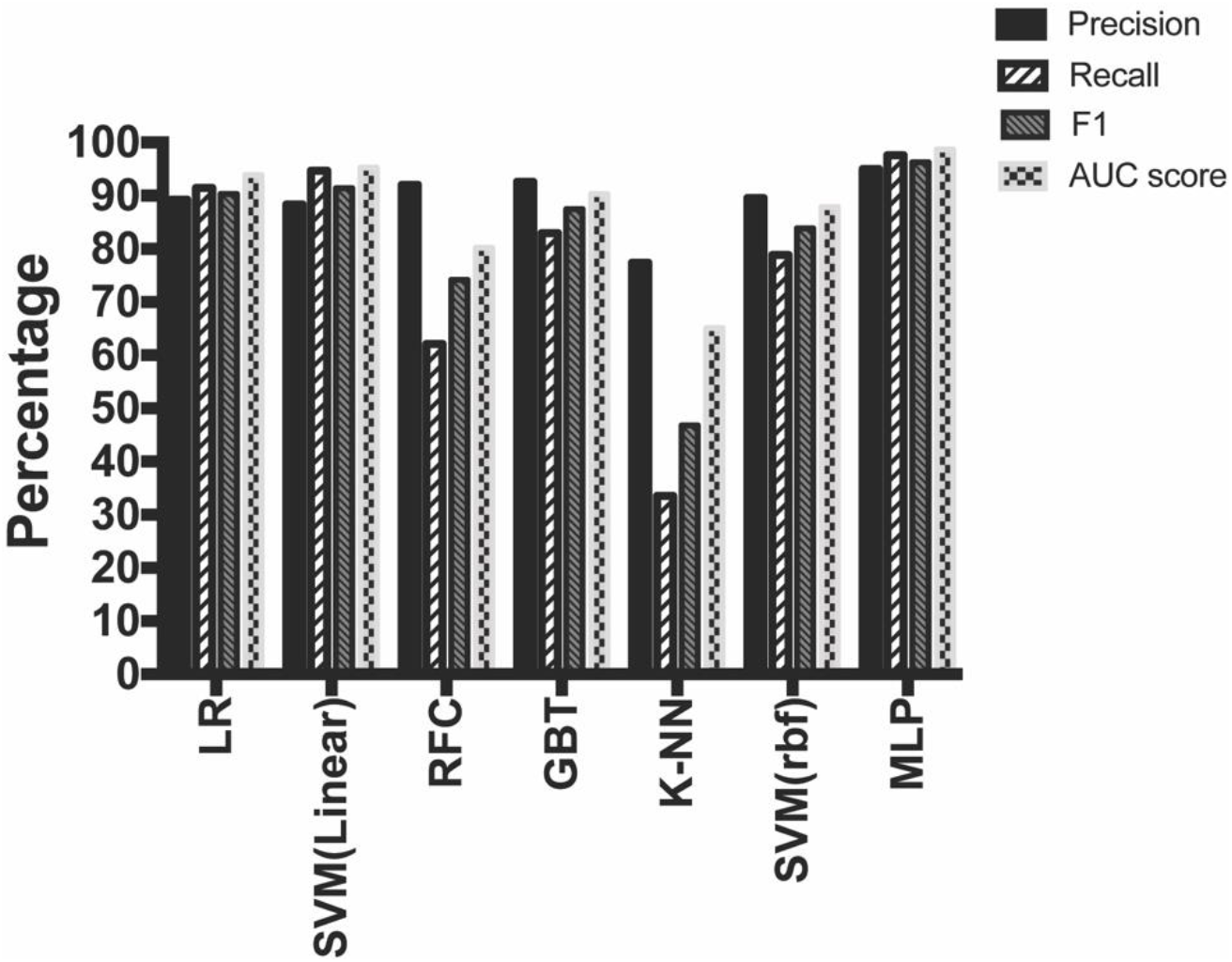
Performance of different machine learning models in the holdout test. Penalized logistic regression (LR); linear SVM; random forest classifier (RFC); gradient boosted tree (GBT); k-nearest neighbor algorithm (K-NN); SVM with rbf kernel; multiple layer perceptron (MLP).

With the parameters tuned, the models were evaluated within the holdout test. The machining learning models achieved different performances in the somatic mutation prediction (Fig. S6). The MLP model achieved the highest precision (95%) in holdout test, and was selected as our predictive model for somatic mutation prediction (Table S3). In short, MLP is a class of feedforward artificial neural network. The tuned MLP model has two layers and each layer with 10 and 5 neurons respectively. Learning rate was set as ‘invscaling’ and sover was set as ‘lbfgs’. The ‘logistic’ activation function was applied in the MLP model.

**Table S3:** The number of mutations in the train set and test set and the prediction performance achieved in the multiple layer perceptron (MLP). The germline variants were treated as predictive negative and somatic mutations were treated as predictive positive. Using tuned multiple layer perceptron model to predict mutations in the holdout test set, the AUC score, precision, recall, and F-Measure are reported.

|  |  | Train Set | Test Set |
| --- | --- | --- | --- |
|  | Germline | 10464 | 3315 |
|  | Somatic | 2817 | 1115 |
| **MLP** |  |  |  |
| AUC | Precision | Recall | F1-score |
| 0.98 | 0.95 | 0.98 | 0.96 |

### Validate predicted somatic mutations

We applied the tuned MLP model to predict germline variants/somatic mutations on the mutations derived from the178 benign biopsies without matching germline DNA. In total, out of the 93,653 mutations, 38,210 were predicted to be somatic mutations. To estimate the overall accuracy of predicted somatic mutations, we randomly selected and genotyped three samples for an evaluation study. Three samples were separately LCM dissected. The extracted DNA were genotyped using Infinium Exome-24 Kit. Genotyped probes with GCSCORE larger than 0.15 from the three samples were mapped to hg19 assemblies. Overlapped coordinates were retrieved and the allele frequencies derived from both technologies were compared. When the allele frequency difference was larger than a certain threshold, we considered the somatic mutation call as false positive. With different allele frequency cutoffs, we plotted the accuracy rate in Fig. S7. When the cutoff is 25% (half of 50%), in other words, mutation calls with allele frequency difference larger than 25% between the two technologies were considered as wrong calls, the accuracy of our called somatic mutations was 80.00%.

**Fig. S7:**
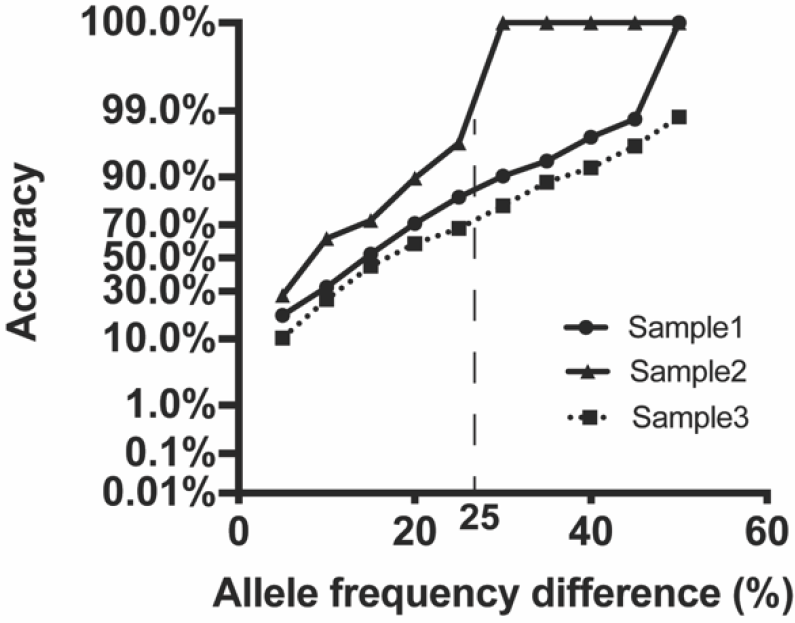
Comparison of the called mutations between genotype array and WES in the cancer samples in the three validation samples. Somatic mutations were called using Mutect2 and our developed somatic mutation prediction model. With different allele frequency difference cutoff, we obtained the accuracy plot.

### Identify somatic mutations in the matched cancer samples

The case group is defined as benign biopsies that developed breast cancer at least one year later after the biopsy. In the case group, we have retrieved 10 matched cancer blocks from the matched individual (Fig. S1). The same procedures were performed as benign biopsies, including LCM dissection, DNA extraction, library construction, sequencing, alignment, mutation calling, and filtering. Specifically, due the relatively higher allele frequency in tumor tissues^30^, the read depth cutoff was adjusted to 10, instead of 20. This is a cutoff widely used in other studies^39^. In total, 10402 mutations were identified in these ten cancer samples.

We then applied ISOWN^82^, which is a predictive model somatic mutation identification in tumors without matched germline DNA. ISOWN utilized ten features for the prediction tasks, include intrinsic and external features. To ensure high precision, we have set stringent criteria and applied naïve Bayes classifier for the prediction. In total, 125 mutations out of 10402 were predicted to be somatic mutations, these mutations distributed in 111 chromosomal locations.

